# powsimR: Power analysis for bulk and single cell RNA-seq experiments

**DOI:** 10.1101/117150

**Authors:** Beate Vieth, Christoph Ziegenhain, Swati Parekh, Wolfgang Enard, Ines Hellmann

## Abstract

Power analysis is essential to optimize the design of RNA-seq experiments and to assess and compare the power to detect differentially expressed genes in RNA-seq data. PowsimR is a flexible tool to simulate and evaluate differential expression from bulk and especially single-cell RNA-seq data making it suitable for a priori and posterior power analyses.

## Introduction

RNA-sequencing (RNA-seq) is an established method to quantify levels of gene expression genome-wide [17]. Furthermore, the recent development of very sensitive RNA-seq protocols, such as Smart-seq2 and CEL-seq [7,18] allows transcriptional profiling at single-cell resolution and droplet devices make single cell transcriptomics high-throughput, allowing to characterize thousands or even millions of single cells [9,15, 30].

Even though technical possibilities are vast, scarcity of sample material and financial consideration are still limiting factors [31], so that a rigorous assessment of experimental design remains a necessity [1, 3]. The number of replicates required to achieve the desired statistical power is mainly determined by technical noise and biological variability [3] and both are considerably larger if the biological replicates are single cells. Crucially, it is common that genes are detected in only a subset of cells and such dropout events are thought to be rooted in the stochasticity of single-cell library preparation [8]. Thus dropouts in single-cell RNA-seq are not a pure sampling problem that can be solved by deeper sequencing [2]. In order to model dropout rates it is absolutely necessary to model the mean-variance relationship inherent in RNA-seq data. Even though current power assessment tools use the negative binomial or similar models that have an inherent mean-variance relationship, they do not explicitly estimate and model the observed relationship, but rather draw mean and variance separately (reviewed in [19]).

In powsimR, we have implemented a flexible tool to assess power and sample size requirements for differential expression (DE) analysis of single cell and bulk RNA-seq experiments. For our read count simulations, we (1) reliably model the mean, dispersion and dropout distributions as well as the relationship between those factors from the data. (2) Simulate read counts from the empirical mean-variance- and dropout relations, while offering flexible choices of the number of differentially expressed genes, effect sizes and DE testing method. (3) Finally, we evaluate the power over various sample sizes. We will use the embryonic stem cell data from [10] to illustrate powsimR’s utility to plan and evaluate RNA-seq experiments.

## powsimR

### Estimation of RNA-seq Characteristics

An important step in the simulation framework is the reliable representation of the characteristics of the observed data. In agreement with others [5,14,16], we find that the read distribution for most genes is sufficiently captured by the negative binomial. We analyzed 18 single cell datasets using unique molecular identifiers (UMIs) to control for amplification duplicates and 20 without duplicate control. The negative binomial provides an adequate fit for 54% of the genes for the non-UMI-methods and 39% of the genes for UMI-methods, while the zero-inflated negative binomial was only adequate for 2.8% of the non-UMI-methods. In contrast, for the UMI-methods a simple Poisson distribution fits well for some studies [25,31] (Supplementary File S2). Furthermore, when comparing the fit of the other commonly used distributions, the negative binomial was most often the best fitting one for both non-UMI (57%) and UMI-methods (66%), while the zero inflated negative binomial improves the fit for only 19% and 1.6% (Supplementary Figure S4). Therefore the default sampling distribution in powsimR is the negative binomial (Figure 1), however the user has also the option to choose the zero-inflated negative binomial.

**Figure 1.**
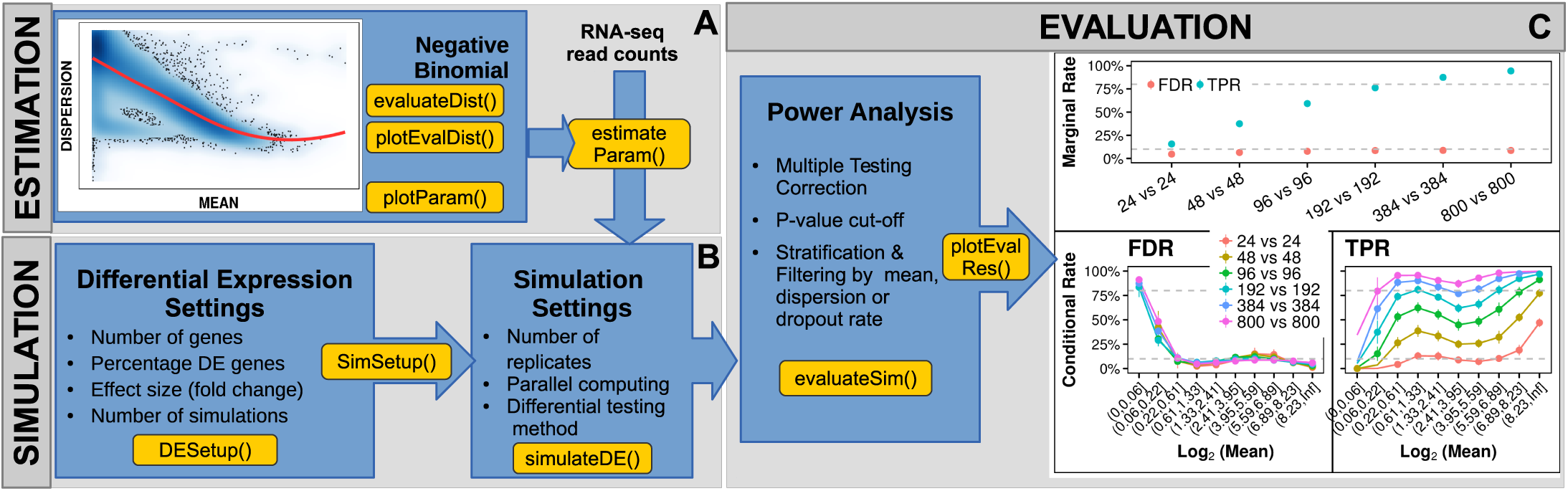
PowsimR schematic overview. **A** The mean-dispersion relationship is estimated from RNA-seq data, which can be either single cell or bulk data. The users can provide their own count tables or one of our five example data sets. The plot shows the mean-dispersion estimated, assuming a negative binomial for the Kolodziejczyk-data, the red line is the loess fit, that we later use for the simulations. **B** These distribution parameters are then used to set-up the simulations. For better comparability, the parameters for the simulation of differential expression are set separately. **C** Finally, the TPR and FDR are calculated. Both can be either returned as marginal estimates per sample configuration (top), or stratified according to the estimates of mean expression, dispersion or dropout-rate (bottom).

### Simulation of Read Counts and Differential Expression

Simulations in powsimR can be based on provided data or on user-specified parameters. We first draw the mean expression for each gene. The expected dispersion given the mean is then determined using a locally weighted polynomial regression fit of the observed mean-dispersion relationship and to capture the variability of the observed dispersion estimates, a local variability prediction band (*σ* = 1.96) is applied to the fit (Figure 1A). Note, that using the fitted mean-dispersion spline is the feature that critically distinguishes powsimR from other simulation tools that draw the dispersion estimate for a gene independently of the mean. Our explicit model of mean and dispersion across genes allows us to reproduce the mean-variance as well as mean-dropout relationship observed (Supplementary Figure S2, Supplementary File S2).

To simulate DE genes, the user can specify the number of genes as well as the fraction of DE genes as *log*_2_ fold changes (LFC). For the Kolodziejczyk data, we found that a narrow gamma distribution mimicked the observed LFC distribution well (Supplementary Figure S3). The set-up for the expression levels and differential expression can be re-used for different simulation instances, allowing an easier comparison of experimental designs.

Finally, the user can specify the number of samples per group as well as their relative sequencing depth and the number of simulations. The simulated count tables are then directly used for DE analysis. In powsimR, we have integrated 8 R-packages for DE analysis for bulk and single cell data (limma [21], edgeR [22], DESeq2 [13], ROTS [24], baySeq [6], DSS [29], NOISeq [26], EBSeq [12]) and five packages that were specifically developed for single-cell RNA-seq (MAST [4], scde [8], BPSC [27], scDD [11], monocle [20]). For a review on choosing an appropriate method for bulk data, we refer to the work of others e.g. [23]. Based on our analysis of the single-cell data from [10], using standard settings for each tool we found that MAST performed best for this dataset given the same simulations as compared to results of other DE-tools.

### Evaluating Statistical Power

Finally, powsimR integrates estimated and simulated expression differences to calculate marginal and conditional error matrices. To calculate these matrices, the user can specify nominal significance levels, methods for multiple testing correction and gene filtering schemes. Amongst the error matrix statistics, the power (True Positive Rate; TPR) and the False Discovery Rate (FDR) are the most informative for questions of experimental design. For easy comparison, powsimR plots power and FDR for a list of sample size choices either conditional on the mean expression [28] or simply as marginal values (Figure 1). For example for the Kolodziejczyk data, 384 single cells for each condition would be sufficient to detect > 80% of the DE genes with a well controlled FDR of 5%. Given the lower sample sizes actually used in [10], our power analysis suggests that only 60% of all DE genes could be detected.

In summary, powsimR can not only estimate sample sizes necessary to achieve a certain power, but also informs about the power to detect DE in a data set at hand. We believe that this type of posterior analysis will become more and more important, if results from different studies are compared. Often enough researchers are left to wonder why there is a lack of overlap in DE-genes when comparing similar experiments. Powsim will allow the researcher to distinguish between actual discrepancies and incongruities due to lack of power.

## Availability

The R package and associated tutorial are freely available at https://github.com/bvieth/powsimR.

## References

1. Paul L Auer and R W Doerge. Statistical design and analysis of RNA sequencing data. Genetics, 185(2):405–416, June 2010.

2. Rhonda Bacher and Christina Kendziorski. Design and computational analysis of single-cell RNA-sequencing experiments. Genome Biol., 17:63, 7 April 2016.

3. Ana Conesa, Pedro Madrigal, Sonia Tarazona, David Gomez-Cabrero, Alejandra Cervera, Andrew McPherson, Micha l Wojciech Szczessniak, Daniel J Gaffney, Laura L Elo, Xuegong Zhang, and Ali Mortazavi. A survey of best practices for RNA-seq data analysis. Genome Biol., 17:13, 26 January 2016.

4. Greg Finak, Andrew McDavid, Masanao Yajima, Jingyuan Deng, Vivian Gersuk, Alex K Shalek, Chloe K Slichter, Hannah W Miller, M Juliana McElrath, Martin Prlic, Peter S Linsley, and Raphael Gottardo. MAST: a flexible statistical framework for assessing transcriptional changes and characterizing heterogeneity in single-cell RNA sequencing data. Genome Biol., 16(1):1–13, 10 December 2015.

5. Dominic Grün, Lennart Kester, and Alexander van Oudenaarden. Validation of noise models for single-cell transcriptomics. Nat. Methods, 11(6):637–640, June 2014.

6. Thomas J Hardcastle. Generalized empirical bayesian methods for discovery of differential data in high-throughput biology. Bioinformatics, 32(2):195–202, 15 January 2016.

7. Tamar Hashimshony, Florian Wagner, Noa Sher, and Itai Yanai. CEL-Seq: single-cell RNA-Seq by multiplexed linear amplification. Cell Rep., 2(3):666–673, 27 September 2012.

8. Peter V Kharchenko, Lev Silberstein, and David T Scadden. Bayesian approach to single-cell differential expression analysis. Nat. Methods, 11(7):740–742, July 2014.

9. Allon M Klein, Linas Mazutis, Ilke Akartuna, Naren Tallapragada, Adrian Veres, Victor Li, Leonid Peshkin, David A Weitz, and Marc W Kirschner. Droplet barcoding for single-cell transcriptomics applied to embryonic stem cells. Cell, 161(5):1187–1201, 21 May 2015.

10. Aleksandra A Kolodziejczyk, Jong Kyoung Kim, Jason C H Tsang, Tomislav Ilicic, Johan Henriksson, Kedar N Natarajan, Alex C Tuck, Xuefei Gao, Marc Bühler, Pentao Liu, John C Marioni, and Sarah A Teichmann. Single cell RNA-Sequencing of pluripotent states unlocks modular transcriptional variation. Cell Stem Cell, 17(4):471–485, 1 October 2015.

11. Keegan D Korthauer, Li-Fang Chu, Michael A Newton, Yuan Li, James Thomson, Ron Stewart, and Christina Kendziorski. A statistical approach for identifying differential distributions in single-cell RNA-seq experiments. Genome Biol., 17(1):222, 25 October 2016.

12. Ning Leng, John A Dawson, James A Thomson, Victor Ruotti, Anna I Rissman, Bart M G Smits, Jill D Haag, Michael N Gould, Ron M Stewart, and Christina Kendziorski. EBSeq: an empirical bayes hierarchical model for inference in RNA-seq experiments. Bioinformatics, 29(8):1035–1043, 15 April 2013.

13. Michael I Love, Wolfgang Huber, and Simon Anders. Moderated estimation of fold change and dispersion for RNA-seq data with DESeq2. Genome Biol., 15(12):550, 2014.

14. Aaron T L Lun, Karsten Bach, and John C Marioni. Pooling across cells to normalize single-cell RNA sequencing data with many zero counts. Genome Biol., 17:75, 27 April 2016.

15. Evan Z Macosko, Anindita Basu, Rahul Satija, James Nemesh, Karthik Shekhar, Melissa Goldman, Itay Tirosh, Allison R Bialas, Nolan Kamitaki, Emily M Martersteck, John J Trombetta, David A Weitz, Joshua R Sanes, Alex K Shalek, Aviv Regev, and Steven A McCarroll. Highly parallel genome-wide expression profiling of individual cells using nanoliter droplets. Cell, 161(5):1202–1214, 21 May 2015.

16. Gu Mi, Yanming Di, and Daniel W Schafer. Goodness-of-fit tests and model diagnostics for negative binomial regression of RNA sequencing data. PLoS One, 10(3):e0119254, 18 March 2015.

17. Ali Mortazavi, Brian A Williams, Kenneth McCue, Lorian Schaeffer, and Barbara Wold. Mapping and quantifying mammalian transcriptomes by RNA-Seq. Nat. Methods, 5(7):621–628, July 2008.

18. Simone Picelli, Omid R Faridani, Asa K Björklund, Gösta Winberg, Sven Sagasser, and Rickard Sandberg. Full-length RNA-seq from single cells using smart-seq2. Nat. Protoc., 9(1):171–181, January 2014.

19. Alicia Poplawski and Harald Binder. Feasibility of sample size calculation for RNA-seq studies. Brief. Bioinform., 18 January 2017.

20. Xiaojie Qiu, Andrew Hill, Jonathan Packer, Dejun Lin, Yi-An Ma, and Cole Trapnell. Single-cell mRNA quantification and differential analysis with census. Nat. Methods, 14(3):309–315, March 2017.

21. Matthew E Ritchie, Belinda Phipson, Di Wu, Yifang Hu, Charity W Law, Wei Shi, and Gordon K Smyth. limma powers differential expression analyses for RNA-sequencing and microarray studies. Nucleic Acids Res., 43(7):e47, 20 April 2015.

22. Mark D Robinson, Davis J McCarthy, and Gordon K Smyth. edger: a bioconductor package for differential expression analysis of digital gene expression data. Bioinformatics, 26(1):139–140, 1 January 2010.

23. Nicholas J Schurch, Pietá Schofield, Marek Gierlinski, Christian Cole, Alexander Sherstnev, Vijender Singh, Nicola Wrobel, Karim Gharbi, Gordon G Simpson, Tom Owen-Hughes, Mark Blaxter, and Geoffrey J Barton. How many biological replicates are needed in an RNA-seq experiment and which differential expression tool should you use? RNA, 28 March 2016.

24. Fatemeh Seyednasrollah, Krista Rantanen, Panu Jaakkola, and Laura L Elo. ROTS: reproducible RNA-seq biomarker detector—prognostic markers for clear cell renal cell cancer. Nucleic Acids Res., page gkv806, 11 August 2015.

25. M Soumillon, D Cacchiarelli, S Semrau, and others. Characterization of directed differentiation by high-throughput single-cell RNA-Seq. bioRxiv, 2014.

26. Sonia Tarazona, Pedro Furio-Tarí, David Turrà, Antonio Di Pietro, María Jos’e Nueda, Alberto Ferrer, and Ana Conesa. Data quality aware analysis of differential expression in RNA-seq with NOISeq R/Bioc package. Nucleic Acids Res., 43(21):e140, 2 December 2015.

27. Trung Nghia Vu, Quin F Wills, Krishna R Kalari, Nifang Niu, Liewei Wang, Mattias Rantalainen, and Yudi Pawitan. Beta-Poisson model for single-cell RNA-seq data analyses. Bioinformatics, 32(14):2128–2135, 15 July 2016.

28. Angela R Wu, Norma F Neff, Tomer Kalisky, Piero Dalerba, Barbara Treutlein, Michael E Rothenberg, Francis M Mburu, Gary L Mantalas, Sopheak Sim, Michael F Clarke, and Stephen R Quake. Quantitative assessment of single-cell RNA-sequencing methods. Nat. Methods, 11(1):41–46, January 2014.

29. Hao Wu, Chi Wang, and Zhijin Wu. A new shrinkage estimator for dispersion improves differential expression detection in RNA-seq data. Biostatistics, 14(2):232–243, April 2013.

30. Grace X Y Zheng, Jessica M Terry, Phillip Belgrader, Paul Ryvkin, Zachary W Bent, Ryan Wilson, Solongo B Ziraldo, Tobias D Wheeler, Geoff P McDermott, Junjie Zhu, Mark T Gregory, Joe Shuga, Luz Montesclaros, Jason G Underwood, Donald A Masquelier, Stefanie Y Nishimura, Michael Schnall-Levin, Paul W Wyatt, Christopher M Hindson, Rajiv Bharadwaj, Alexander Wong, Kevin D Ness, Lan W Beppu, H Joachim Deeg, Christopher McFarland, Keith R Loeb, William J Valente, Nolan G Ericson, Emily A Stevens, Jerald P Radich, Tarjei S Mikkelsen, Benjamin J Hindson, and Jason H Bielas. Massively parallel digital transcriptional profiling of single cells. Nat. Commun., 8:14049, 16 January 2017.

31. Christoph Ziegenhain, Beate Vieth, Swati Parekh, Björn Reinius, Amy Guillaumet-Adkins, Martha Smets, Heinrich Leonhardt, Holger Heyn, Ines Hellmann, and Wolfgang Enard. Comparative analysis of Single-Cell RNA sequencing methods. Mol. Cell, 65(4):631–643.e4, 16 February 2017.

